# A boosted unbiased molecular dynamics method for predicting ligands binding mechanisms: Probing the binding pathway of dasatinib to Src-kinase

**DOI:** 10.1101/650440

**Authors:** Farzin Sohraby, Mostafa Javaheri Moghadam, Masoud Aliyar, Hassan Aryapour

## Abstract

Small molecules such as metabolites and drugs play essential roles in biological processes and pharmaceutical industry. Knowing their interactions with biomacromolecular targets demands a deep understanding of binding mechanisms. Dozens of papers have suggested that discovering of the binding event by means of conventional unbiased molecular dynamics (MD) simulation urges considerable amount of computational resources, therefore, only one who holds a cluster or a supercomputer can afford such extensive simulations. Thus, many researchers who do not own such resources are reluctant to take the benefits of running unbiased molecular dynamics simulation, in full atomistic details, when studying a ligand binding pathway. Many researchers are impelled to be content with biased molecular dynamics simulations which seek its validation due to its intrinsic preconceived framework. In this work, we have presented a workable stratagem to encourage everyone to perform unbiased (unguided) molecular dynamics simulations, in this case a protein-ligand binding process, by typical desktop computers and so achieve valuable results in nanosecond time scale. Here, we have described a dynamical binding’s process of an anticancer drug, the dasatinib, to the c-Src kinase in full atomistic details for the first time, without applying any biasing force or potential which may lead the drug to artificial interactions with the protein. We have attained multiple independent binding events which occurred in the nano-second timescales, surprisingly as little as ∼30 ns. Both the protonated and deprotonated forms of the dasatinib reached the crystallographic binding mode without having any major intermediate state during induction.

## Introduction

Small molecules play essential roles in almost all cellular mechanisms. Studying of how they bind to other macro-molecules and trigger a specific activity can unravel interesting information. Over the last decades, sophisticated methods such as X-ray crystallography, NMR and electron microscopy have revealed numerous structural details of many protein-ligand complexes (ww, 2019) by producing detailed pictures of them. However, these pictures are just one or some static poses of a vivid system which its function can be swayed by its movements and dynamics. Furthermore, in many proteins like androgen receptor (PDB ID: 2Q7I) (Askew, et al., 2007), the binding pocket is buried deep inside the protein structure. The experimental methods cannot reveal much detail about the process of induction and, in a whole picture, the binding pathway of them. How a ligand makes an entrance into the protein and how it affects residues on its journey to reach the native binding pose have been left on the sidelines, most often.

By now, there have been many unbiased molecular dynamics studies which mostly have led us to profound results and subsequently enhanced our understanding about binding mechanisms (Buch, et al., 2011; Dickson and Lotz, 2016; Dickson and Lotz, 2017; Dror, et al., 2011; Gohlke, et al., 2013; Mondal, et al., 2018; Schneider, et al., 2016; Shan, et al., 2011; Stanley, et al., 2016). Unfortunately, these studies are computationally very expensive, and the implementaion of unbiased MD is limited to few institutions who can afford supercomputers. Dror et al., for example, performed about 270 microseconds of unbiased MD simulations on β2-adrenergic receptor. Out of 82 independent simulations, each lasting from 1 to 19 microseconds, they eventually reached 21 binding events (Dror, et al., 2011). Stanley et al. resolved the binding of a lipid inhibitor, the ML056, to the sphingosine-1-phosphate receptor 1 after 800 microseconds, utilizing unbiased MD (Stanley, et al., 2016). Shan et al. also resolved the bindings of the PP1 and the dasatinib to the c-Src kinase protein by using unbiased MD, with a total simulation time of 115 and 35 microseconds respectively (Shan, et al., 2011). These large-time scales may discourage many scientists to perform unbiased MD simulations, obviously, because they may think this method is very pricy. Recently made breakthroughs in the context of biased methods have permitted determination of some binding kinetics of a protein-ligand complex (Casasnovas, et al., 2017; Decherchi, et al., 2015; Guo, et al., 2016; Limongelli, et al., 2013; Motta, et al., 2018; Sinko, et al., 2013; Soldner, et al., 2019; Tiwary, et al., 2015; Tiwary, et al., 2017; You and Chang, 2018). Although these advancements have enhanced sampling by utilizing either modification of both potential energy surface and temperature or reassignment of probability weight factor, they mainly have ended up either in uncertainty in total potential energy or substandard depiction of canonical ensemble. Besides, employment of biased methods requires precise selection of biasing variables and potentials, a crucial step that any mistakes in it will lead to false positive results.

Nevertheless, we believe this dilemma can be addressed by Unaggregated Unbiased Molecular Dynamics (UUMD) simulation. In this study, we have presented new information about the sampling, induction and binding of the dasatinib in complex with the c-Src protein kinase by utilizing the UUMD which means we have avoided to apply any biasing forces or potentials between the dasatinib and the protein. Our team decided to pick up the dasatinib/c-Src kinase complex, because the unbiased MD simulation of this complex had been conducted formerly by Shan et al. (Shan, et al., 2011), so we would be able to set their work as a benchmark for our study. Make it possible to be employed by any researchers, we have tried using the unbiased MD in our own way. We gained interesting structural details of the binding process of the dasatinib in complex with the c-Src kinase that had not been published before.

## Methods

### Construction of systems

The X-ray crystallography of human c-Src kinase proteins with the PDB ID: 1Y57 (Cowan-Jacob, et al., 2005) was obtained from the Protein Data Bank. Missed side chains and atoms were added and refined using the UCSF Chimera (Pettersen, et al., 2004). Then, co-crystallized ligands and water molecules were removed. The remainder was the apo-protein structure which after some picoseconds of equilibration was ready for further unbiased MD simulations. Hetero ligands (protonated and deprotonated dasatinibs) were parameterized, using the default settings of the ACEPYPE (Sousa da Silva and Vranken, 2012) for assigning the partial charges and atom types. The ionization state of the dasatinib is in the pH range 6-8 with the pKa around 7.29 (Gaulton, et al., 2017). Hence, in the physiological pH, there can be seen two possible forms of the dasatinib, protonated and deprotonated. Because both forms can be presented inside cells, we took both of them into account in our simulations. It is noteworthy to say existence of a positive formal charge at the tail segment of dasatinib - all segments will be defined further – can overshadow the entire characteristic of the compound. For determining binding pathway of the dasatinib, a high concentration of this ligand and c-Src kinase was inserted into the simulation box. The first round made up of several short-run simulations, instead of one long-run simulation, which differed only based on their initial velocities and constituted both protonation states. Utilizing these short runs, we were able to single out some configurations which had the potentials to bind successfully. The key criteria for the selection of these likely to bind configurations were their orientations’ favorability and their conformations’ closeness to the X-ray structure (PDB ID: 3G5D) (Getlik, et al., 2009). Further details are presented by a flowchart (Figure 2).

**Figure 1.**
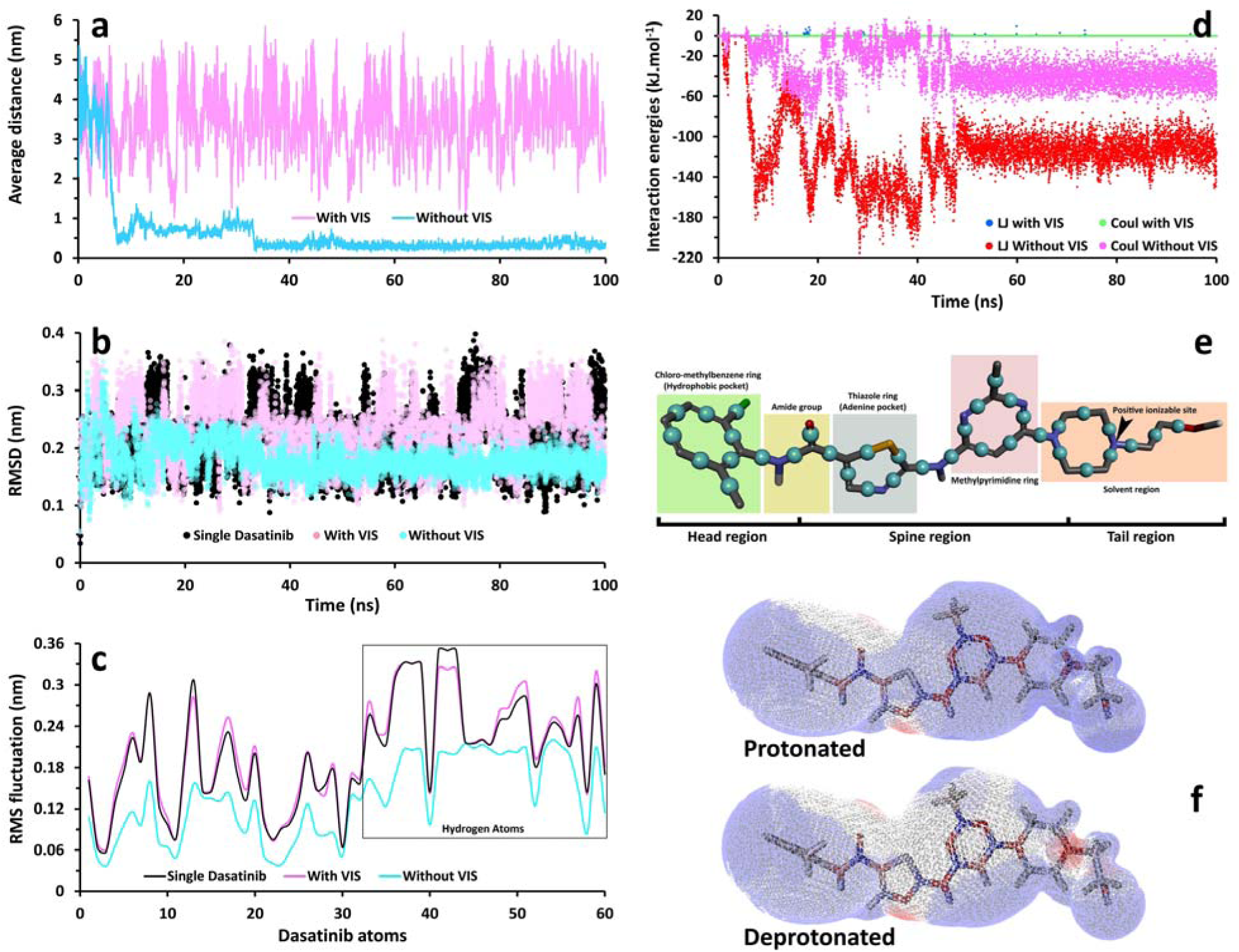
The influence of virtual interaction sites (VIS) on molecular behavior of the dasatinib in the simulation box which held 16 of them. **a**, The average distance between two randomly selected dasatinib molecules once in the presence of VISs and once in the absence of them. **b**, Changes of the RMSD values, regarding two types of the dasatinib molecules; those provided with VISs (VIS-equipped) and those remained untouched (VIS-unequipped). In both case the molecules were selected randomly. The outputs, then, were compared to another single dasatinib molecule which was simulated individually (control plot). All RMSDs were calculated based on the initial conformations of the related dasatinib molecules. After 35 ns, the RMSD values of the VIS-unequipped dasatinib molecules fell down, and it fluctuated between 0.1 and 0.25 nm onwards. In contrast, in the presence of VISs the RMSD values were the subject of more fluctuations, in the range of 0.1 to 0.4 nm, which was similar to the control plot. **c**, Root mean square fluctuation (RMSF) of the VIS-unequipped dasatinibs. The fluctuation of the VIS-equipped dasatinibs was almost as same as the control plot. On the contrary, in the absence of VIS the fluctuation of dasatinibs decreased significantly. **d**, The Van der Waals (VdW) and Coulombic (Coul) interactions energies between the VIS-equipped dasatinibs and the VIS-unequipped dasatinibs. In the absence of VIS, dasatinibs aggregated and their behavior were under the influence of some strong attractive non-covalent forces (especially Van der Waals). The employment of the repulsive force evened up the attractive force. **e**, The fragmented form of the dasatinib structure, the VISs are represented by dark cyan balls placed between heavy atoms of the dasatinib to distribute repulsive potential evenly all over the molecule. **f**, The comparison of partial charge surfaces in the two forms of dasatinib molecule. Atoms are colored based on their partial charges; red, blue and grey colors indicate negative, positive and neutral charges, resp.

**Figure 2.**
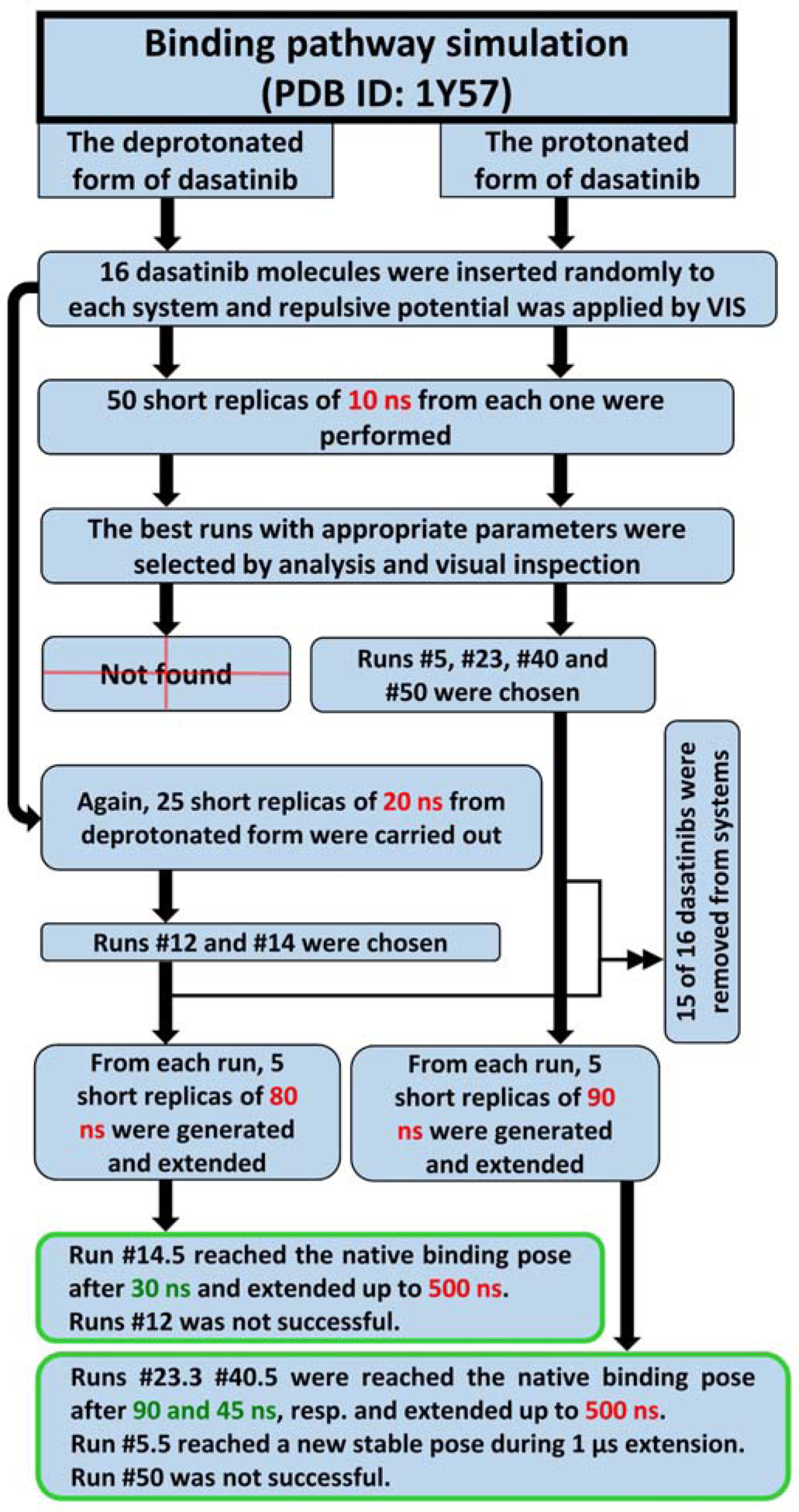
Unaggregated unbiased molecular dynamics flowchart. All replicas were identical except in their velocities.

### MD simulation protocol and analyses

All simulations were initiated without or with pre-equilibrated state of the relevant apo-protein using the OPLS force field (Kaminski, et al., 2001) in GROMACS 2018 (Abraham, et al., 2015), respectively. There were 100 simulation boxes in total, 50 for each ionization states of the dasatinib. In each triclinic box, apart fromTIP3P waters (Jorgensen, et al., 1983), there were 16 ligands (the dasatinib) which were randomly positioned as well as the apo-protein (c-Src kinase) which was placed at the center of the box, with a distance of 1.5 nm from all edges. To neutralize boxes and also satisfy physiological salt concentration of 150 mM sodium and chloride ions were added. Final number of atoms in each system reached up to 42000. Then each system went through energy minimization, using steepest descent algorithm, until the Fmax met below 10 kJ.mol^-1^.nm^-1^. All bonds were constrained using the Linear Constraint Solver (LINCS) algorithm (Hess, et al., 1997). The long-range electrostatic interactions were treated, using the Particle Mesh Ewald (PME) method (Darden, et al., 1993), and the cut off radii for Coulomb and Van der Waals short-range interactions was set to 0.9 nm. To keep the system in the stable environmental conditions (310 K, 1 Bar), the modified Berendsen (V-rescale) thermostat (Bussi, et al., 2007) and Parrinello–Rahman barostat (Parrinello and Rahman, 1981) were applied for 100 and 300 ps respectively. Finally, simulations were carried out under the periodic boundary conditions (PBC). The subsequent analyses were then performed using GROMACS utilities, VMD (Humphrey, et al., 1996) and USCF Chimera. The associated plots were produced, implementing Daniel’s XL Toolbox (v 7.3.2) add-in (Kraus, 2014). The free energy landscapes were rendered, utilizing Matplotlib (Hunter, 2007). In addition, we estimated the binding free energy by taking the advantage of the g_mmpbsa package (Kumari, et al., 2014). All of the computations were performed on an Ubuntu desktop PC, with a [dual-socket Intel(R) Xeon(R) CPU E5-2630 v3 + 2 NVIDIA GeForce GTX 1080] configuration. The approximate performance of this platform is ∼250 ns/day if running a system which has ∼40000-atom.

## Results

### The optimization of unbiased MD simulation

Usually, small molecules tend to aggregate together to form a ball-like structure; even two ligands can aggregate inside a simulation box for an extensive amount of time. Dwindling sampling effort, this can increase the required simulation time substantially. On the other hand, sampling by only one ligand, demands lots of computational time. Adaption of repulsive potential among ligands can be a promising stratagem in this respect. While putting this potential into practice can prohibit ligands’ aggregation, it has no significant impact on other interactions. Shan et al. applied a weak repulsive force among “the nitrogen atoms of the central amide group of dasatinib molecules”, with a cut off of 2 nm. They reached one successful binding event after a total run time of 32 µs, by inserting 6 molecules inside the simulation box. Apart from this work, by now there have been many other studies that have employed unbiased MD simulation to reconstruct the binding event (Dror, et al., 2011; Mondal, et al., 2018; Schneider, et al., 2016; Stanley, et al., 2016) but none of these studies have yet levied a repulsive force among ligands. In fact, most of them either used only one ligand or went through consecutive ligands’ aggregations during their simulations, especially when the ligands’ concentration was high. Furthermore, the aggregation can limit the ligands’ movements in the solvent and, therefore, alter their natural behavior. In our UUMD simulations, we applied repulsive forces between Virtual Interaction Sites (VIS) which were placed evenly all over the molecule, almost between each two heavy atoms of the ligand (Fig. 1 e). The repulsive force was formulated through the reassignment of the Lennard-Jones’ parameters (Fig. 1d). The values of the σ and ε parameters for VISs were set to 0.83 nm and 0.1 kJ.mol^-1^, respectively. While our applied repulsive force prevents ligands to adsorb to each other, it has almost no impact on the native fluctuations and behavior of the ligands, even at considerably high concentrations. The results show that all of the 16 inserted ligands in the simulation box acted as similar as a single solvated ligand, even when the concentrations were as high as 60 mM (Fig. 1a, b, c). This approach allowed us to enhance the chance of complete sampling and so the probability of achieving a successful binding event, without dealing with some undesirable changes in protein structure (Ext Fig. 2). A series of short runs (nanosecond time scale simulations) were carried out, instead of conducting one long run (microsecond time scale simulation) (Fig. 2) to foster the sampling’s process of the protein’s surface, even more.

In the first attempt, 100 independent simulations were conducted, 50 for each of ionization states of dasatinib. All simulations started with identical atomic coordinates, but different velocities, called replicas. Time length of each replica did not exceed of 10 ns and each held 16 dasatinib molecules. For protonated form of dasatinib, after careful analyses of the first round of simulations, 4 runs (#5, #23, #40 and #50) in which dasatinibs were sufficiently well-oriented, considering their conformations’ closeness to the co-crystallized dasatinib, were singled out and pipelined to the next round of simulations. At this point, in each of these four simulation boxes only one dasatinib molecule which was highlighted as a well-oriented molecule during previous step was kept. The remainder dasatinib molecules were removed. Next, 5 replicas were generated from each of these 4 boxes, 20 runs in total, and then simulations were resumed from where it was ended and continued up to 100 ns. Out of these 20 runs, two attempts, #23.3 and #40.5, demonstrated successful inductions due to the RMSD criteria. Their RMSD values reached less than 2 Å, regarding the crystal structure as the reference structure. Then each of these two runs, #23.3 and #40.5, were taken up again; each extended up to 500 ns. Nonetheless, a new binding pose was also found out. Out of five replicas of run #5, one (run #5.5) showed a new binding pose. It was then extended up to 1 µs (Ext Fig. 1). Short-run stratagem has been also examined by Knapp et al. where they confirmed that short replicas are far more efficient than long-time runs in terms of protein folding (Knapp, et al., 2018).

Sampling of the binding pocket was more challenging when the deprotonated form of dasatinib was employed. In the first round of simulations, no successful run has been found. Thus, the second round of simulations was launched, 25 replicas at durations of 20 ns. As the result, two attempts (#12 and #14) were detected. They were presenting well-orientated dasatinibs which were good candidates for further simulations. So, from each of these two runs 5 replicas at durations of 80 ns were made. Finally, in one replica (#14.5) dasatinib reached the native binding pose (the crystal structure). By extending this run up to 500 ns, its placement was confirmed.

Surprisingly, by taking this approach the chance of good sampling, especially around the binding site, can be increased significantly and in its wake achieving native binding poses would be much more frequent and quicker. By applying the present method, it took just 30 ns for the ligand (dasatinib) to reach its native binding pose while it took 2.3 µs for Shan et al.

### The study of the binding pathways of dasatinib

As far as we know, no detailed report about how the dasatinib binds to the c-Src kinase has been published by now. By analyzing all of the simulated trajectories, we unravel some of fine characteristics of the protein structure and also describe how and in what different modes dasatinib can bind to this protein. The c-Src kinase domain has two main parts (Fig. 3a); a big carboxyl-terminus lobe and a small amino-terminus lobe. The binding pocket exists in the middle of these two lobes which forms a cleft. These two main parts are joined by the αC-helix and a very important random-coil structure which is called “the hinge” (Roskoski, 2015).

**Figure 3.**
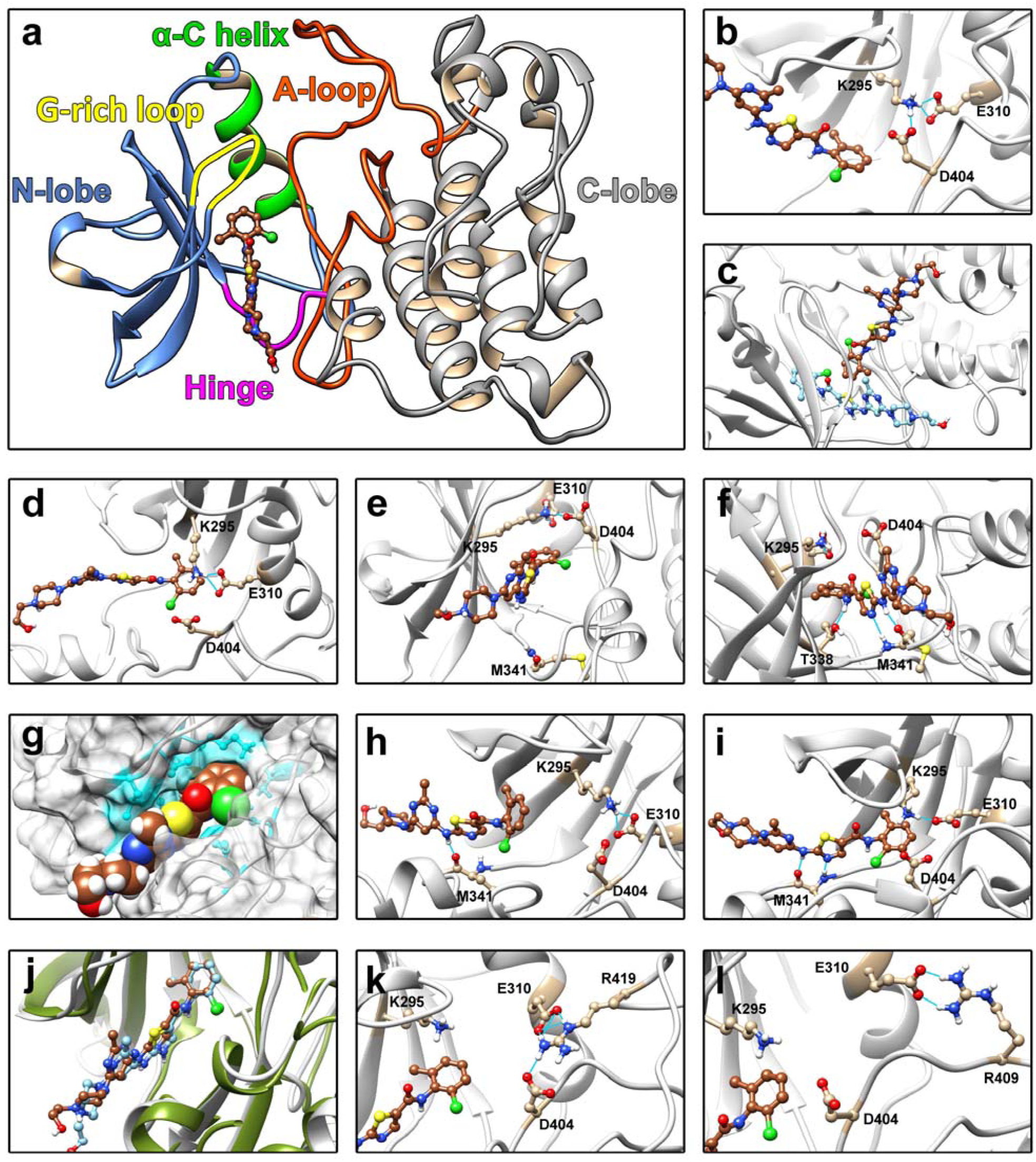
The atomic details of the dasatinib binding pathways. **a**, The entire structure of c-SRC kinase domain with its key regions. **b**, Before the induction, K295, E310 and D404 get together and forms the salt bridges. **c**, The well-oriented dasatinib. The head pointed down towards the deep parts of the binding pocket. The X-ray structure of dasatinib is shown in cyan. Further selections (runs #5, #12, #14, #23, #40 and #50) were picked up based on this orientation. **d**, The disruption of the salt bridges by the head of dasatinib. As it can be seen, It has broken the salt bridge between K295 and D404, but the other one which was established between K295 and E310, was stronger, and so remained unbroken. **e**, Just before the landing of the spine of dasatinib on the hinge. **f**, The landing of the spine and in turn formation of two hydrogen bonds between the backbone of M341 and the two Nitrogen atoms of dasatinib. The formation of another hydrogen bond, between T338 and the head of dasatinib, is also illustrated. **g**, The hydrophobic interactions between the dasatinib, presented at both the spine and the head segments, and the hydrophobic residues of the protein, including L273, V281 and L393 which embraced the dasatinib. They are colored in cyan. **h**, The second binding mode; Only one hydrogen bond between the oxygen atom of the backbone of M341 and the dasatinib was formed, obviously in the first step. **i**, Then, the head of dasatinib went down inside the binding pocket and got underneath the conserved salt bridges. A second hydrogen bond was formed between T338 and the amide group of the dasatinib. **j**, The native binding pose achieved by UUMD simulation. **K**, The interference of the A-loop. The R419 broke the conserved salt bridges and confiscated both of the E310 and D404. **l**, Formation of another salt bridge between the R409 and E310 took place when the glutamate residue had been headed outwards. This could result in the αC-helix-out conformation.

Before dasatinib induces the binding pocket, deep parts of the binding pocket are inaccessible, mainly due to the presence of two evolutionary conserved salt bridges which were established among three residues (K295, E310 and D404). These salt bridges are, in fact, the indicators of the active form of the protein (Meng and Roux, 2014) (Fig. 3b). In addition, existence of some hydrophobic residues on both sides of the cleft, interacting with each other time to time, close the binding pocket occasionally. In the two attempts, #23.3 and #40.5, dasatinib induced the binding pocket when its head segment had been orientated towards the deep parts of the binding pocket (Fig. 3c). As it was going down inside the pocket, its hydrophobic interactions, especially at its hefty chlorine substitute, with the protein got stronger and the head of dasatinib weakened the salt bridges. In other words, once the salt bridges were weakened, the head of dasatinib went deeper into the binding pocket quickly (Fig. 3d). Then, the spine segment of dasatinib landed on the hinge part and subsequently two hydrogen bonds were formed between the backbone of M341 and the two nitrogen atoms, located at the spine of dasatinib (Fig. 3e, f). Based on the interaction energies, the M341 has the most contribution in the stability of the dasatinib (Ext Fig. 3). These two aforementioned hydrogen bonds stimulated the hydrophobic interactions between the spine of dasatinib and the hydrophobic residues like the L273, V281, L393 and Y340. These residues, in turn, embraced the dasatinib and so preserved it from surrounded water molecules (Fig. 3g). In row, the tail segment of dasatinib was being almost solvated.

In spite of the discussed binding mode, a second binding pathway was also observed in the run #14.5. Although binding elements had not been changed, the sequence of the incidents was different. In this run, the spine of dasatinib was the first region which landed on the hinge and then only one hydrogen bond was formed between the oxygen atom of the M341 and the dasatinib. From this point and onwards, the hydrophobic interactions were stabling the dasatinib (Fig. 3h). Next, the head of dasatinib made its way into the deep parts of the binding pocket, underneath the conserved salt bridges. Simultaneously, a second hydrogen bond was established between the nitrogen atoms of the M341 and the dasatinib, provoked the native binding-pose (Fig. 3i). The calculated values of K_on_ for the both forms of the dasatinib were equal to 7.56 s^-^ 1.µM^-1^, compare to the experimentally determined value (5 s^-1^.µM^-1^) (Shan, et al., 2009).

The head of dasatinib requires a considerable amount of space and when this fragment is in the deep parts of the binding pocket, it can weaken the evolutionary conserved salt bridges, established among three residues K295, E310 and D404. In other words, the presence of the dasatinib inside the pocket is in row with the probability of the salt bridges’ break-off. Interestingly, these salt bridges somehow play as the stabilizers of dasatinib’s head segment. While side chains of the K295 and the E310 were, frequently, making hydrophobic and coulombic interactions with the head of dasatinib, their absence made the dasatinib more unstable (Fig. 4a). Breakage of these salt bridges can pave the way for the nearby water molecules to flow inside the binding pocket and keep remained in deep parts of this pocket for a considerable amount of time. This can also cause higher rate of fluctuations in both of the K295 and the E310 which in its wake can make the head of dasatinib more unstable.

**Figure 4.**
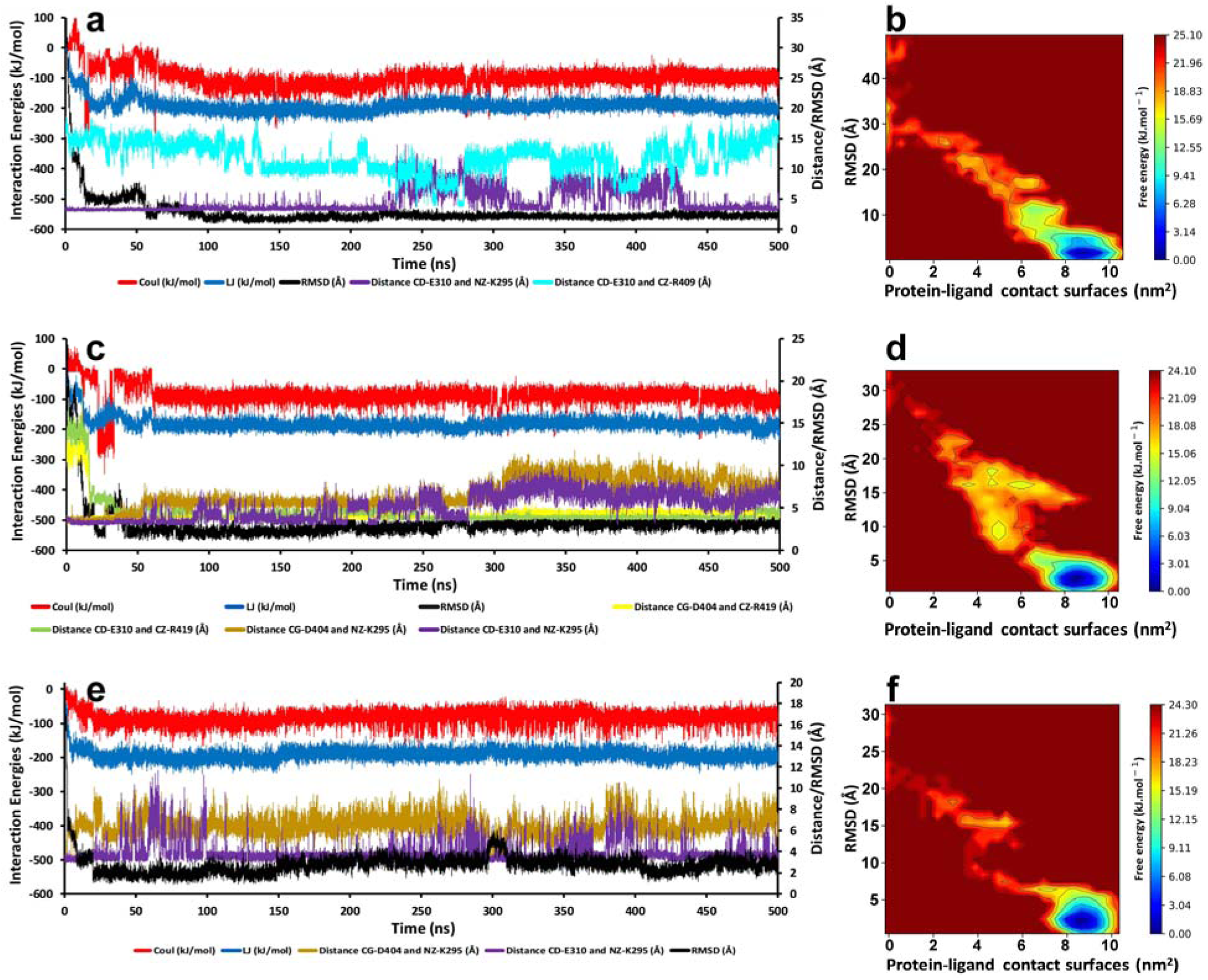
Energy figures as well as the RMSD values regarding the binding pathway of the protonated and deprotonated forms of the dasatinib to the Src-kinase. **a**, Run #23.3. Atomic distance between CD of E310 and NZ of K295, represents the breakage and then reformation of this salt bridge throughout the simulation time. The K295 and D404 did not make any salt bridge during this run. The RMSD values as well as interaction energies of the protein-ligand complex converged at 0.9 Å just after 90 ns and fluctuated between 1 and 2 Å onwards. From 230 to 450 ns, the salt bridges were broken so E310 was able to make a bond with R409 which resulted to instability of the dasatinib’s head. This can be seen through a slight rise in both of the interaction energies and the RMSD values. **b**, The Free Energy Landscape of the run #23.3 which was calculated by using “gmx sham”. Shows that in the binding process no intermediate states were made and the binding happened in a short amount of time. **c**, Run #40.5. Dasatinib reached the native binding pose in just 45 ns. The atomic distances between K295 and either of E310/D404 (salt bridges) as well as between R419 and either of E310/D404 (salt bridges) show that from 20 ns the R419, by breaking the anterior salt bridges, started to confiscate both of D404 and E310 and had been preserving this situation by the end of time frame. This opens the way for dasatinib to bind. **d**, The free energy landscape of the run #40.5. **e**, Run #14.5. Dasatinib reached the native binding pose in just 30 ns. **f**, The free energy landscape of the run #14.5.

The breakage of the salt bridges which triggers a cascade of significant conformational changes in the protein structure (McClendon, et al., 2014; Roskoski, 2015), can take place under the influence of three different factors; (i) Kinetic energies of bonded residues and water molecules. Take the periodic motions of the E310, from the inner side of the pocket towards the surface and its interactions with water molecules is a good illustration of this. (ii) Mechanical energy, derived from the head fragment of dasatinib. (iii) Activation loop (A-loop). There are two key arginine residues in the A-loop, the R419 and R409, which can interfere with the above-described salt bridges. The R419 can bind to the both of D404 and E310 at the same time (Fig. 3k). In the run #40.5, we found that whenever the R419 confiscates these two acidic residues, dasatinib, consequently, can reach to the deep parts of the binding pocket much easier (Fig. 4c). The R409 was also able to bind to the E310 while it was heading outwards. This incident could result in the αC-helix-out conformation (Lin, et al., 2013; McClendon, et al., 2014; Shukla, et al., 2014) (Fig. 4a, Fig. 3j). It was also observed that these two arginine residues could even swap E310 with each other. Generally, the A-loop can bind to the revolutionary conserved E310 and D404, and transform active conformation of the protein into inactive form, utilizing R419 and R409. It was previously reported that only the R409 could trigger this conformational change (Shukla, et al., 2014), but we found the R419 can also play a significant role. These conformational changes are probably under the A-loop’s control which itself undergoes major conformational changes throughout this transformation.

The T338 pulls the head of dasatinib toward itself by establishing an influential hydrogen bond with the amide group, near the head of dasatinib, so that water molecules cannot seep into this area. However, this hydrogen bond can be easily broken-off by either water mediation or the rotation of the T338 side-chain’s dihedral angle. Although the methyl group of T338 can form Alkyl-Pi interaction with the head of dasatinib, its impact on the captivation of the dasatinib’s head is not as substantial as the hydrogen bond.

In this work, UUMD simulation enabled us to reveal different probable mechanisms for the binding of dasatinib to the c-Src kinase. Moreover, this method is even capable of predicting binding conformations and binding pockets that cannot be achieved by the x-ray crystallography (data are presented in the supplementary material file, run #5.5).

## Conclusion

Herein, we have introduced an approach to encourage other researchers to do unbiased MD simulations of binding events with a reasonable computational resource. By imposing repulsive forces among the ligands and also utilization of rational sampling the binding process were noticeably speeded up. While the sampling of the target protein was enhanced by introducing high concentrations of the ligands, the efficiency of this process was also increased, utilizing short runs, instead of one long run. The calculated K_on_ was fairly close to the experimental one which can confirm our approach. In addition, this approach can help researchers to spot out new binding pockets as well as native binding poses, and also in terms of drug resistance and optimization studies.

In drug discovery projects, on both commercial and academic grounds, efficiency is of the immense significance. We have achieved this great efficiency by employing the OPLS force field. Nevertheless, other force fields may be more efficient and challenging as well. This question can be answered in future works. Screening, identification and optimization of potential compounds based on the target macromolecule structure can be considered as some other appliances of the UUMD. It could also help to solve many challenges ahead of docking algorithms including binding pocket identification, ligand sampling, protein flexibility, and scoring functions.

## Supporting information

Run #5.5

Run #14.5

Run #23.3

Run #40.5

## Acknowledgment

This work was supported by the grant number 96-1206 from Golestan University, Gorgan, Iran.

## Conflict of interest

The authors declare no conflict of interest.

## Notes

### Competing Interest Statement

The authors have declared no competing interest.

### Summary of Updates

Dear Editor; The main text and its contents have been revised. Also, Supplemental files revised and reuploaded. To avoid occupying more space, you could delete previous video links and replace them with new ones. Sincerely, Hassan Aryapour

