## Supplementary materials for "A boosted unbiased molecular dynamics method for predicting ligands binding mechanisms: Probing the binding pathway of dasatinib to Src-kinase"

**Results**

In the run #5.5, we witnessed an unusual binding conformation which had not been reported by anyone before. We found that the protonated form of dasatinib could bind in a 180-degrees flipped conformation, in comparison to the native binding pose reported by the x-ray crystallography (Ext Fig. 1e).

At the first step, the R419 triggered the process by confiscating the E310 and D404. This was accompanied by breakage of salt bridges, established among the K295, D404 and E310, and thereafter expansion of the pocket’s room (Ext Fig. 1b). This breakage made the way for F278 to turn inside the binding pocket and stabilize dasatinib by the π-stacking interactions which eventually steered dasatinib to slip deeper into the pocket (Ext Fig. 1c, d). Hydrophobic residues around the pocket also played a supportive role to stabilize this conformation (Ext Fig. 1e). The accumulation of these forces lowered the RMSD values to just above 3 Å.

Although the Van der Waals interaction energy scheme of this new binding pose (#5.5) is very similar to the VdW energy scheme of the native binding poses (#23.3, #40.5 and #14.5), its electrostatic interaction energy scheme was modestly different from the native pose. The electrostatic energy plot for the new binding pose is much more erratic due to the periodic formation and breakage of a salt bridge, made between the tail of dasatinib and D348. It is also evident that the overall electrostatic energy’s trend is more positive in comparison with the native poses (Ext Fig. 1a). However, since, during a micro-second simulation, it was observed that dasatinib could interact with the outer residues which were located around the binding pocket and been pulled out occasionally, this binding pose may not be as stable as the native conformation (Ext Fig. 1f).

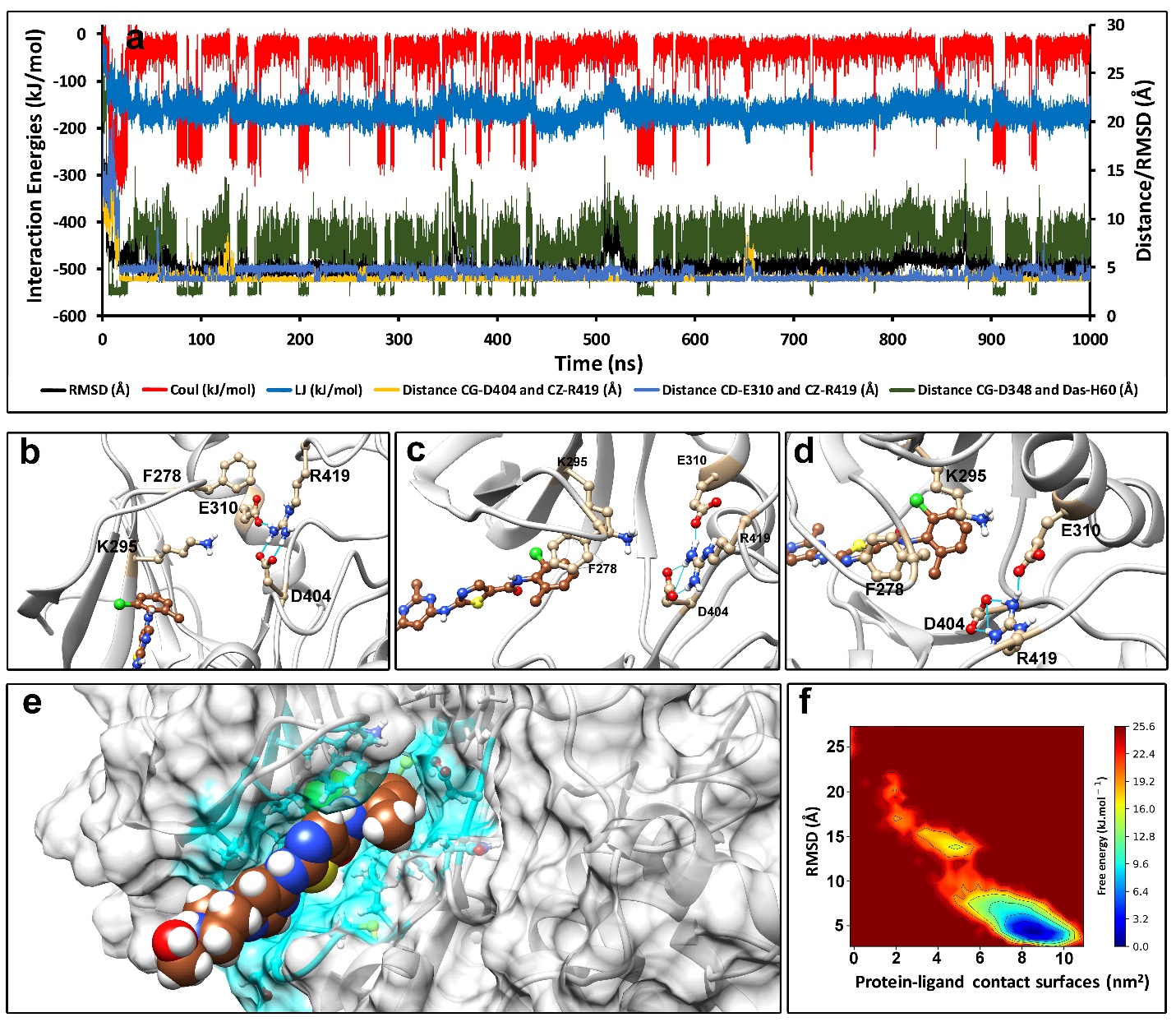

**Ext Fig. 1. A overall scheme of the new binding mode for dasatinib in the run #5.5. a,** The interaction energies of the protein-ligand complex and the changes of distances between the R419 and either of D404/E310 during 500 ns simulation. Soon after the simulation was begun, the R419 started to confiscate both of E310 and D404, and then had been preserving this conformation until the end of simulation. The Van der Waals interaction energy was relatively similar to the native bindings, but the electrostatic energy periodically dropped down in the wake of a salt bridge formation and breakage between the tail of dasatinib and D348. Surprisingly, the RMSD for this novel binding mode reached to 3 Å. **b**, Formation of salt bridge between R419 and both of E310 and D404. **c,** In this new binding mode, the contribution of F278 was detrimental. The F278, situated in the G-rich loop, can turn inside the binding pocket and stabilize the head of dasatinib by a π- stacking interaction. **d,** Dasatinib could get deep inside the binding pocket with the help of F278 and K295. **e,** The surface view of c-SRC kinase in complex with dasatinib. The run #5.5 shed a new light on our understanding about possible binding modes of dasatinib to the c-Src kinase by illustrating a new binding mode which had not been reported before. We found that dasatinib could bind to the c-SRC kinase in a 180-degrees flipped conformation, in comparison to the native binding mode in reference structures (the reported X-ray crystallography structures). **f,** The free energy landscape of dasatinib.

Although for the dasatinib-c-Src kinase complex, we have performed 50 replicas each with a duration of 10 ns, but it should be kept in mind that the duration and the number of the replicas heavily depends on the systems’ characteristics. That is to say, time scales and number of replicas will need to be modified if any component of the system changes. One should set them to suit their system well. In another study, that will be published in the near future, exploring binding mechanism of HTLV1 protease inhibitors, we reached the crystallographic structure, using a protocol which consisted of 100 replicas (each with a duration time of 50 ns). Because the binding site of the inhibitor was covered by two flaps which must be open, in order for the inhibitor to reach inside. Briefly, the duration time and the number of replicas are not something fixed. They mostly depend on structure and complexity of target proteins. The setup must be changed until it satisfies our desired goal. As a good illustration of this in the first round of sampling runs, regarding the deprotonated dasatinib, no runs out of 50 replicas, each with 10 ns duration, was satisfying. As the result, we changed the setup to 25 replicas each with 20 ns of duration time which led to better result.

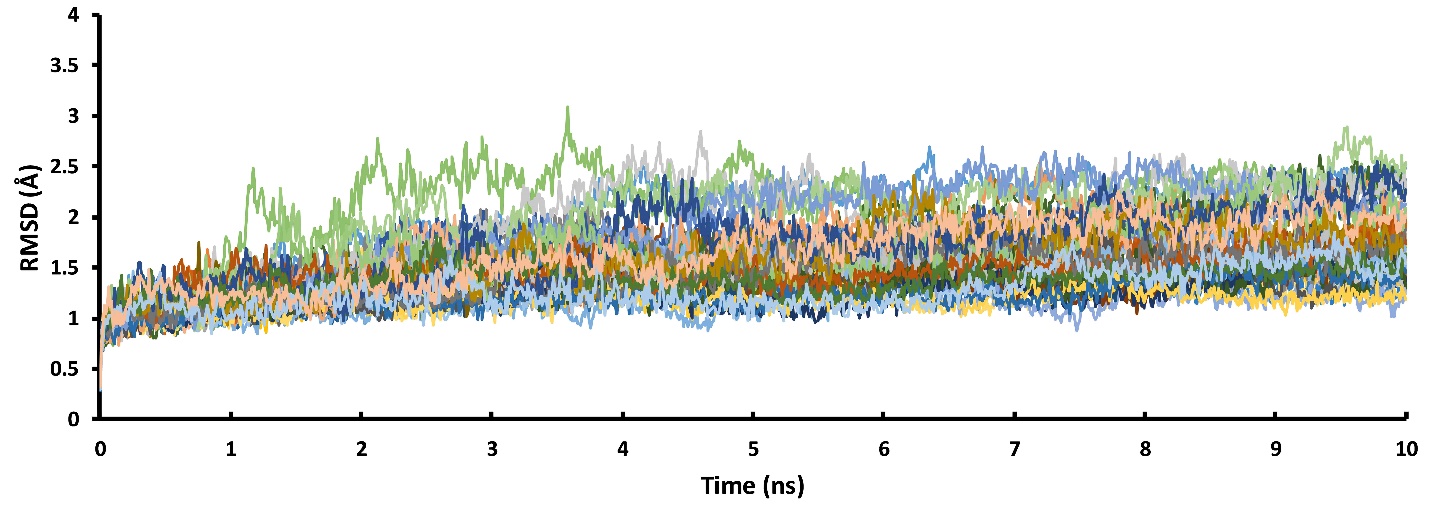

**Ext Fig. 2. The RMSD values of the protein backbone in the 50 sampling runs with 16 ligands present in the simulation box.** The RMSD values show that the high concentration of the ligand has no effect on the folding of the protein.

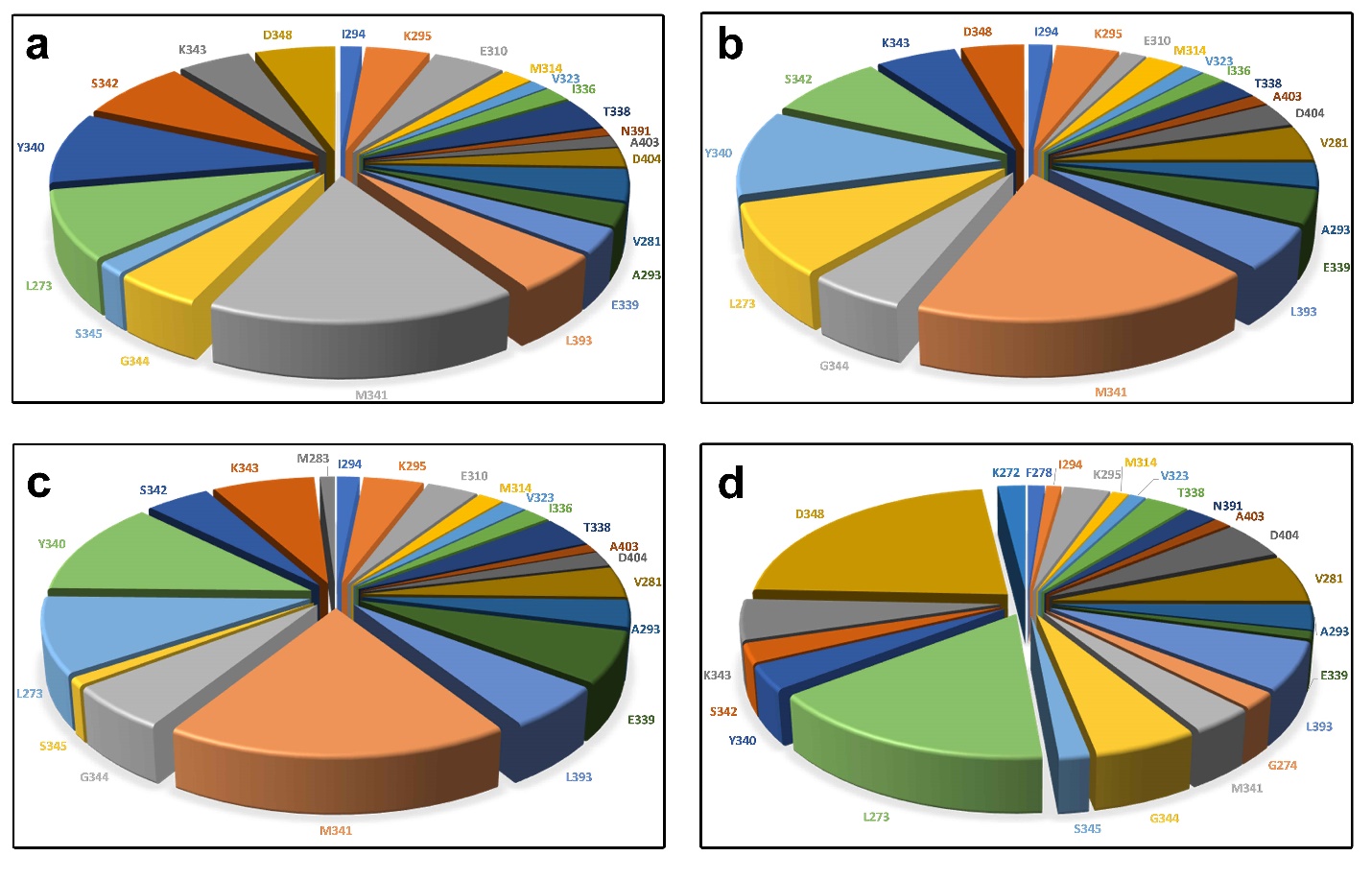

**Ext Fig. 3. The interaction energy contribution of each residue in the protein-ligand binding pathway. a,** run #23.2. **b,** run #40.5. **c,** run #14.5. **d,** run #5.5.
